# Plasma neutralizing antibodies in an infant with interclade HIV-1 superinfection preferentially neutralize superinfecting HIV-1 strains

**DOI:** 10.1101/2020.12.10.420703

**Authors:** Nitesh Mishra, Shaifali Sharma, Ayushman Dobhal, Sanjeev Kumar, Himanshi Chawla, Swarandeep Singh, Ravinder Singh, Bimal Kumar Das, Rakesh Lodha, Sushil Kumar Kabra, Kalpana Luthra

## Abstract

HIV-1 superinfection is defined as infection by an unrelated second strain of HIV-1 after seroconversion due to primary infecting strain and has been associated with development of breadth in the neutralizing antibody (nAb) response, altered disease progression and efficacy of antiretrovirals; though conflicting observations have also been reported. Superinfection has been reported in HIV-1 infected adults. Recently we observed that multivariant infection in infants was associated with early induction of plasma broadly neutralizing antibodies (bnAbs) targeting diverse autologous viruses, however, there is paucity of information on infants with HIV-1 superinfection. Furthermore, the mechanisms by which superinfection in an infant, after priming by an initial infection, potentiate the evolution of a bnAb response have not been evaluated. Herein, we performed a longitudinal analysis and observed evolution of nAb responses in an antiretroviral naïve perinatally HIV-1 infected infant, with interclade superinfection (clade C followed by a unique A1C recombinant). The nAb responses broadened rapidly after superinfection targeted an undefined glycan-dependent epitope on the superinfecting variant, while no enrichment of nAb response against the primary infecting strain occurred. Defining virological features in infants with sequential infection with highly divergent circulating viruses that improve nAb responses will contribute information that could be leveraged for optimization of multicomponent candidate vaccines.

**Importance:** HIV-1 infected infants develop bnAbs rapidly suggesting factors governing bnAb induction in infants are distinct from adults. HIV-1 superinfection is more common in adults whereas the stringent genetic bottleneck for transmission in infants often leads to infection by a single transmitted/founder HIV-1 strain. Longitudinal studies in infants with HIV-1 superinfection can provide key information on the viral factors that induce a bnAb response towards development of a polyvalent vaccine. Herein, we show that in infant who was sequentially infected with two HIV-1 strains from different clades, antibody responses were primarily generated against the superinfecting, second strain of HIV-1.

These antibody responses were dependent on glycans, and targeted an undefined epitope in the C3V4 region of HIV-1 Env. A better understanding of how neutralizing antibody responses develop during natural HIV-1 superinfection in infants will provide information relevant to HIV Env vaccine development and evaluation.

## Introduction

Though a prolonged exposure to viral antigens has been implicated as a necessity for the induction of bnAbs in HIV-1 infected adults, potent plasma bnAbs have been found in infected infants as early as one-year post-infection (1–6). Further, the bnAbs isolated from infected children show features atypical of adult bnAbs suggesting the factors governing bnAb induction in infants are different than those in adults, providing an impetus to better characterize pediatric HIV-1 humoral immunity (7, 8). In a recently reported cohort of 51 HIV-1 infected infants recruited in acute/early stages of infection and screened for plasma neutralization activity against a global panel of twelve heterologous viruses, we had identified a 54-weeks old perinatally HIV-1 infected infant AIIMS709 with two highly divergent and unlinked HIV-1 isolates (24% genetic difference in the env gene between the two contemporaneous circulating viral isolates which is analogous to the difference observed between two distinct clades) who was presumed to be superinfected (5). Using longitudinal samples available from AIIMS709, the current study was designed to address how superinfection in an infant would drive bnAb responses, in the setting of an immature immune system. Considering that the pathways for elicitation of bnAb lineages are suggested to vary significantly between children and adults, we examined the features of the nAb responses in this infant with superinfection and whether the evolving nAbs were targeted at epitopes conserved on both infecting strains or preferentially neutralize either the primary or superinfecting strains.

## Results and Discussion

Two plasma samples from AIIMS709, corresponding to 9-weeks (9-wk) and 54-weeks (54-wk) post detection of HIV-1 infection, prior to initiation of antiretrovirals, were analyzed. To resolve the dynamics of circulating viral populations and multiplicity of infection, we generated a total of 75 single-genome amplified (SGA) env sequences from plasma viral RNA. More than 30 SGA amplicons from each time point were sequenced, providing a 90% confidence interval of sequencing all major viral variants circulating at a population frequency of >5% (**figure 1**). All the viral sequences from the 9-wk plasma sample were closely related, with minimum genetic divergence and formed one defined cluster, as expected, considering the sequences were from an acute stage of infection. Viral env sequences from the 54-wk plasma sample were highly divergent and formed four distinct clusters on the phylogeny tree. Among the 54-wk viral env sequences, cluster 1 sequences were closely related to the 9-wk cluster 1 sequences. The 54-wk clusters 2, 3 and 4 comprised of sequences from circulating viral variants that were highly divergent from 9-wk as well 54-wk cluster 1 sequences and formed a distinct branch on the phylogeny tree (**figure 1**). The env genetic distance between the four viral clusters from 54-wk ranged from 12 to 25%, consistent with unlinked viruses. The intra-cluster evolutionary divergence, as anticipated, was maximal in cluster 3, given cluster 3 variants were the major circulating variants (30 of the 41 SGA env sequences from 54-wk were in cluster 3). All the viruses in 9-wk and 54-wk cluster 1 were classified as clade C while all viral sequences in 54-wk cluster 2, 3 and 4 were unique recombinants between clade A1 and C (A1C) and computationally predicted to be CCR5 tropic. Based on hamming distance frequency distribution of the 54-wk A1C viral variants, superinfection was predicted to have occurred between 24-wk to 30-wk post-detection of HIV-1 infection. Taken together, the sequence analysis performed was suggestive of an initial infection with clade C followed by superinfection with a distinct A1C recombinant HIV-1 variant.

**Figure 1.**
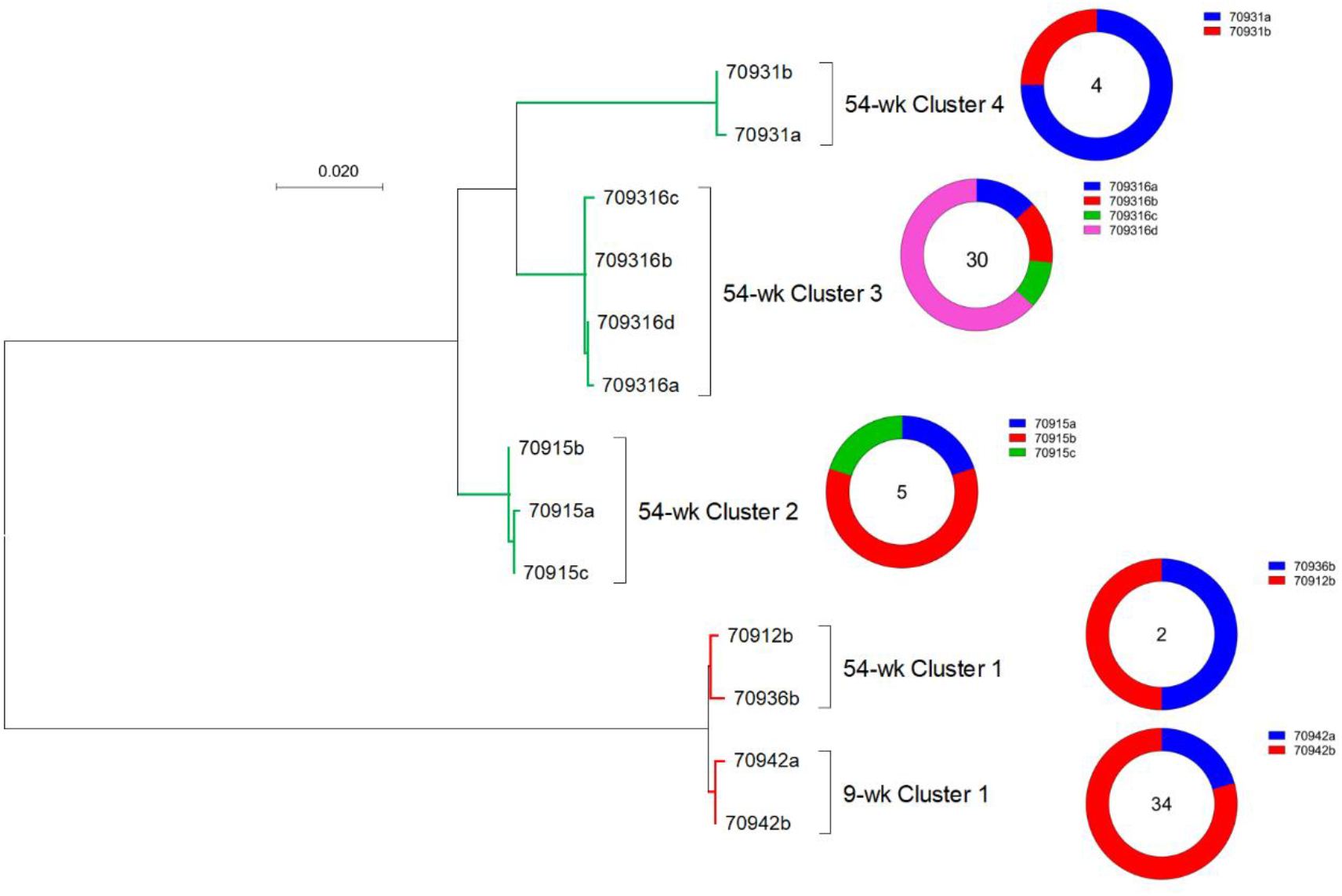
AIIMS709 had interclade superinfection. Maximum-likelihood phylogeny tree of full-length env SGA amplicons (HXB2 position 6225 – 8795) from AIIMS709 9-wk and 54-wk post-detection of HIV-1 infection. Clade C sequences are colored red while A1C variants are colored green. Donut plot adjacent to each cluster represents the number of sequences each branch represents. 54-wk cluster 3 variants were the dominant circulating strains at 54-wk time-point.

We next assessed the HIV-1 specific nAb responses in this superinfected infant from both the available plasma samples (9-wk and 54-wk) to understand the development of nAb breadth, against a panel of 30 pseudoviruses representing major circulating clades and CRFs. While the 9-wk plasma antibodies failed to neutralize majority of the 30-virus panel and only showed strain-specific responses (neutralized 3 clade C viruses of the 30 viruses with ID_50_>50), the 54-wk plasma antibodies neutralized 26/30 viruses (cross-clade neutralization), with a breadth of 86% and a median geometric mean titre (GMT) of 162, (**figure 2A**) which is typical of reported infant plasma nAbs (3, 5, 9). Moreover, the 54-wk plasma nAbs preferentially neutralized clade C viruses showing enrichment of clade specific nAb responses (**figure 2B**). However, considering our 30-virus multiclade panel had relative abundance of clade C viruses (13/30), it may have plausibly skewed the neutralization breadth towards clade C. On a normalized tier scale where plasma nAbs are ranked based on logistic regression to give a tier-like score (10), AIIMS709 54-wk plasma had a score of 3.80 (**figure 2C**), highlighting the ability of AIIMS709 54-wk plasma bnAbs to neutralize majority of the globally circulating tier 3 HIV-1 isolates, albeit with moderate potency.

**Figure 2.**
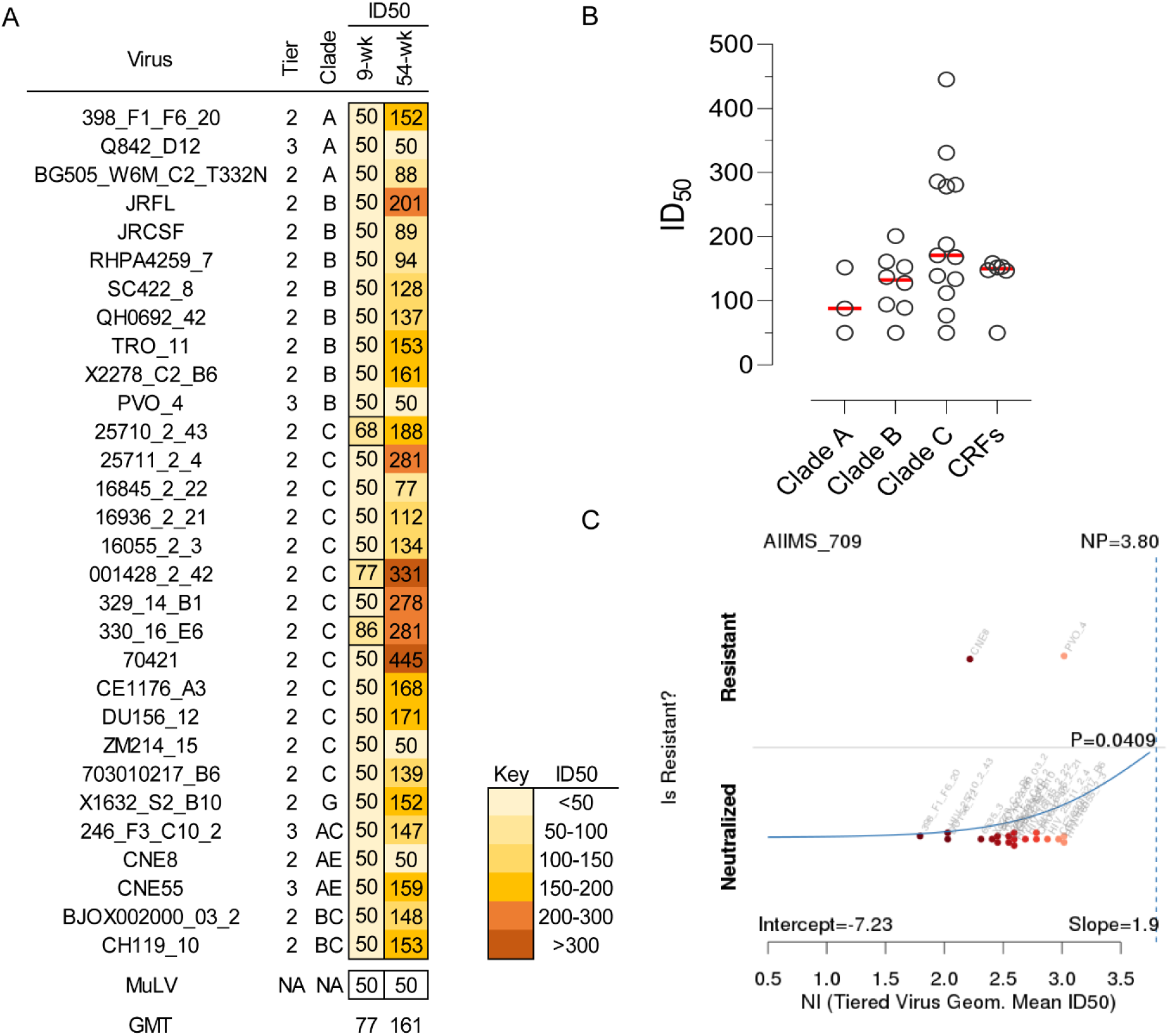
Strain-specific neutralization responses were enriched in AIIMS709. (A) Heatmap representing HIV-1 specific neutralization titres (inverse plasma dilution) of nAbs from AIIMS709 9-wk and 54-wk plasma against the 30-virus multiclade panel. ID_50_ values are color-coded per the given key with darker colors implying higher ID_50_ titres and stronger neutralization. Neutralization assay were repeated thrice in triplicates and average values were used to draw curves and calculate ID_50_ titres. (B) 54-wk ID_50_ titres against individual viruses from the 30-virus panel were grouped according to the clade (A, B, C and CRFs). Median ID50 titres are highlighted as bold red lines. No statistical significance was seen but a trend was observed where clade C viruses had higher ID_50_ titres. (C) Neutralization indexing for AIIMS709 against the 30-virus multiclade panel showed that plasma nAbs were able to neutralize majority of tier 2 viruses on a tier scale that ranks neutralization on a continuous rather than categorical scale.

To define the epitopes targeted by AIIMS709 plasma nAbs, we further employed mutant pseudoviruses containing key mutations in known bnAb epitopes or altered glycan profile, as well as genetic signature analysis associated with neutralization breadth. We did not perform epitope mapping with the 9-wk plasma as it exhibited weak nAb responses. No dominant epitope specificity could be deciphered for the AIIMS709 54-wk plasma nAbs, when tested with mutant pseudoviruses (**figure 3A – E**). The hyperglycosylated variants of 25710_2_43, 16055_2_3, CAP45_G3 and BG505_W6M_C2 pseudoviruses (kifunensine treated mannan-rich and swainsonine treated complex-glycan rich) showed enhanced susceptibility to neutralization by the AIIMS709 plasma bnAbs (**figure 4**), suggesting the presence of glycan-targeting plasma bnAbs in AIIMS709. To confirm that the enhancement in neutralization observed due to hyperglycosylation was not restricted to a single viral backbone, we generated hyperglycosylated variants of all 30-viruses of the multiclade panel tested herein. An overall similar profile of neutralization enhancement of hyperglycosylated viruses was observed, with a 2 – 6-fold increase in ID_50_ titres of plasma bnAbs, showing a preference for mannan-rich glycan than complex glycans (**figure 2b**). Of note, in addition to enhancing neutralization of viruses that were already neutralization sensitive, viruses that were initially resistant to neutralization (Q842_D12, PVO_4, ZM214_15 and CNE8) became susceptible to AIIMS709 plasma nAbs, in their hyperglycosylated form. Intact glycan shields with high glycan density on the viral env have been shown to be enriched for mannans, that in close proximity plausibly sterically restrict the access to mannosidases, resulting in under processing of glycans, thereby leading to recognition by several nAbs (11). Given the established features of several classes of bnAbs that require carbohydrates as either stable anchors or contacts and the improvement seen in neutralizing activity of AIIMS 709 plasma antibodies against hyperglycosylated viruses, we hypothesized that glycan directed nAbs are enriched in the AIIMS709 plasma that can be confirmed by isolation and detailed characterization of the nAbs from this infant (12–14). Computational mapping analysis to predict genetic signatures associated with plasma neutralization breadth suggested AIIMS709 plasma nAbs were dependent on several residues within the C3V4 region of gp120 (**figure 5**). The C3V4 region has been reported as a major nAb target in early subtype C infection, presumably due to its high immunogenicity as a result of increased exposure, being a component of the CD4-binding site as well as the presence of several conserved glycans such as N339, N355 and N363 in the C3 region and N393 and N398 in the V4 region of the viral envelope glycoprotein (15–18). Overall, our findings suggest the presence of plasma nAbs in AIIMS709, of undescribed specificity that are glycan dependent and plausibly target the C3V4 region and can neutralize the hyperglycosylated Env. Next, we generated functional pseudoviruses from the SGA amplicons from each cluster (9-wk and 54-wk) with maximum identity to the cluster consensus sequence and evaluated their susceptibility to neutralization by the autologous 54-wk plasma nAbs. These Env pseudoviruses enabled us to decipher whether the interclade superinfection in this infant promoted the development of cross-clade nAb responses that target the epitopes present on both infecting strains (primary and superinfecting) or are specific to the primary or superinfecting viruses. The 54-wk plasma nAbs showed considerable neutralization of the contemporaneous 54-wk A1C superinfecting viral variants with ID_50_ titres that were more than 25-fold higher compared to the GMT across the multiclade 30-virus panel (ID_50_ of 3149, 3246 and 2854 for 54-wk cluster 2, 3 and 4 viral strains respectively).The 9-wk and 54-wk cluster 1 clade C primary infecting strain showed limited susceptibility to neutralization, as compared to the A1C variants, by the 54-wk plasma nAbs (ID_50_ titres of 348 and 243 respectively) (**figure 6A**). No discernible neutralization of the 9-wk or 54-wk variants with the 9-wk plasma antibodies was observed (data not shown). Considering that the 54-wk plasma nAbs had preferentially neutralized hyperglycosylated heterologous viral strains, we further tested the neutralization susceptibility of the hyperglycosylated autologous viral strains by the plasma nAbs. As observed with heterologous viruses, the hyperglycosylated autologous viruses were better neutralized by AIIMS709 54-wk plasma nAbs. Moreover, in consonance with the observation with hyperglycosylated heterologous viruses, the autologous hyperglycosylated A1C variants showed higher susceptibility to neutralization than that of the clade C hyperglycosylated variants (**figure 6B**). Furthermore, the AIIMS709 54-wk plasma nAbs showed preferential binding to surface expressed envelope glycoprotein from 54-wk cluster 2, 3 and 4 than 9-wk and 54-wk cluster 1 viruses, affirming the preferential recognition of A1C variants by the 54-wk plasma nAbs (**figure 6B**).

**Figure 3.**
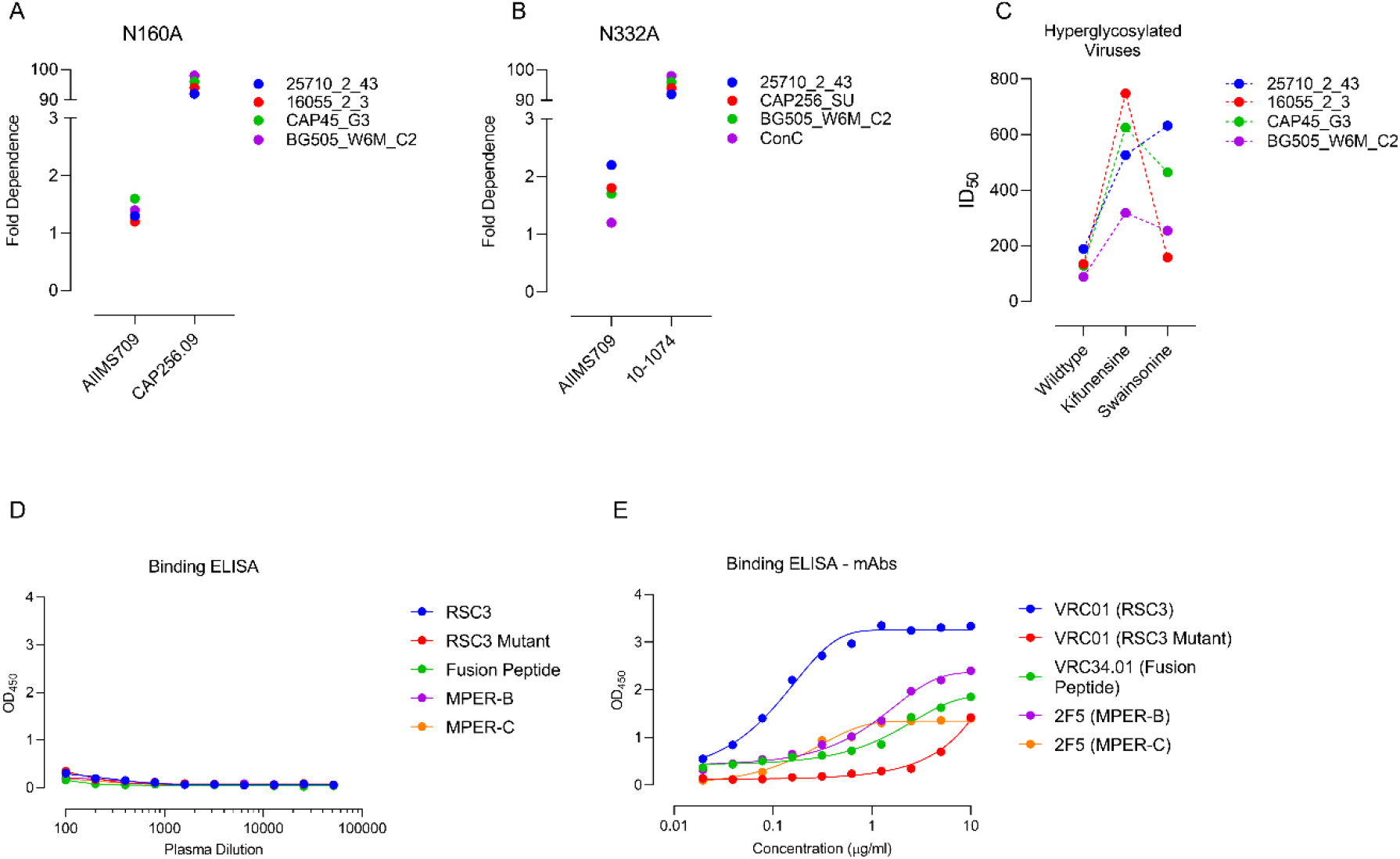
AIIMS709 plasma nAbs target an undefined glycan dependent epitope on HIV-1 Env. (A – B) For V2-apex (N160A) and V3-glycan (N332A) epitope dependence, single base mutants of four HIV-1 pseudoviruses (25710_2_43, 16055_2_3, CAP45_G3 and BG505_W6M_C2 for V2-apex, 25710_2_43, CAP256_SU, BG505_W6M_C2 and ConC for V3-glycan) were used. PG9 (for V2-apex) and 10-1074 (for V3-glycan) were used as positive control. Fold dependence was calculated based on change in ID_50_ against wild type pseudovirus compared to mutant pseudovirus. (C) Comparison of ID_50_ values for 25710_2_43, 16055_2_3, CAP45_G3 and BG505_W6M_C2 and their hyperglycosylated variants grown in presence of kifunensine (ER α – mannosidase I inhibitor) and swainsonine (Golgi α – mannosidase II inhibitor) showed AIIMS709 plasma nAbs prominently neutralized hyperglycosylated viruses. (D) Binding ELISA for CD4bs (RSC3 and RSC3 mutant (Δ371I-P363N)), Fusion peptide and MPER-B and C peptides was performed for AIIMS709 plasma nAbs. OD_450_ values were used to create binding curves. (E) VRC01 (for CD4bs), VRC34.01 (for Fusion peptide) and 2F5 (for MPER-B and -C peptide) were respectively used as positive controls in binding ELISA.

**Figure 4.**
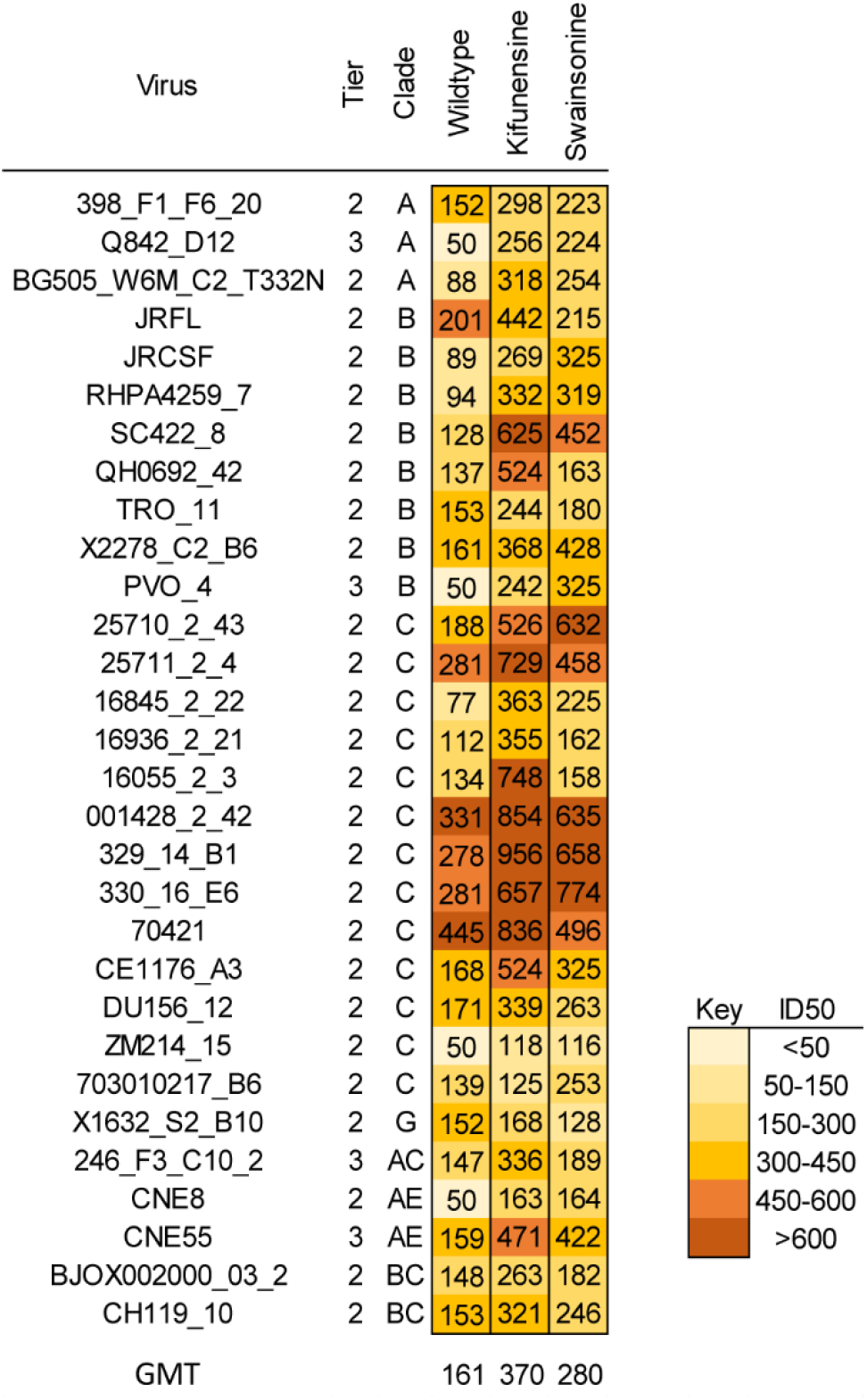
Hyperglycosylation made resistant viruses sensitive to neutralization by AIIMS709 plasma nAbs. Heatmap representing HIV-1 specific neutralization titres (inverse plasma dilution) of nAbs from AIIMS709 54-wk plasma against the 30-virus multiclade panel and their hyperglycosylated variants grown in presence of kifunensine (ER α – mannosidase I inhibitor) and swainsonine (Golgi α – mannosidase II inhibitor). ID_50_ values are color-coded per the given key with darker colors implying higher ID_50_ titres and stronger neutralization. Neutralization assay were repeated thrice in triplicates and average values were used to draw curves and calculate ID_50_ titres.

**Figure 5.**
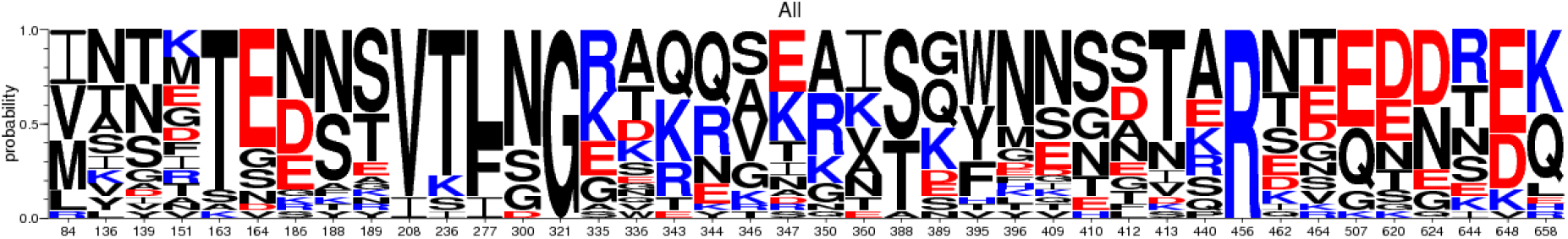
Amino acid signatures associated with neutralization by AIIMS709 plasma nAbs. Amino acid residues are color-coded according to their polarity. Letter height represents the amino acid frequency in the 30-virus multiclade panel. Several residues within the C3V4 region (HXB2 amino acid numbering 332 – 418) showed phylogenetically corrected association.

**Figure 6.**
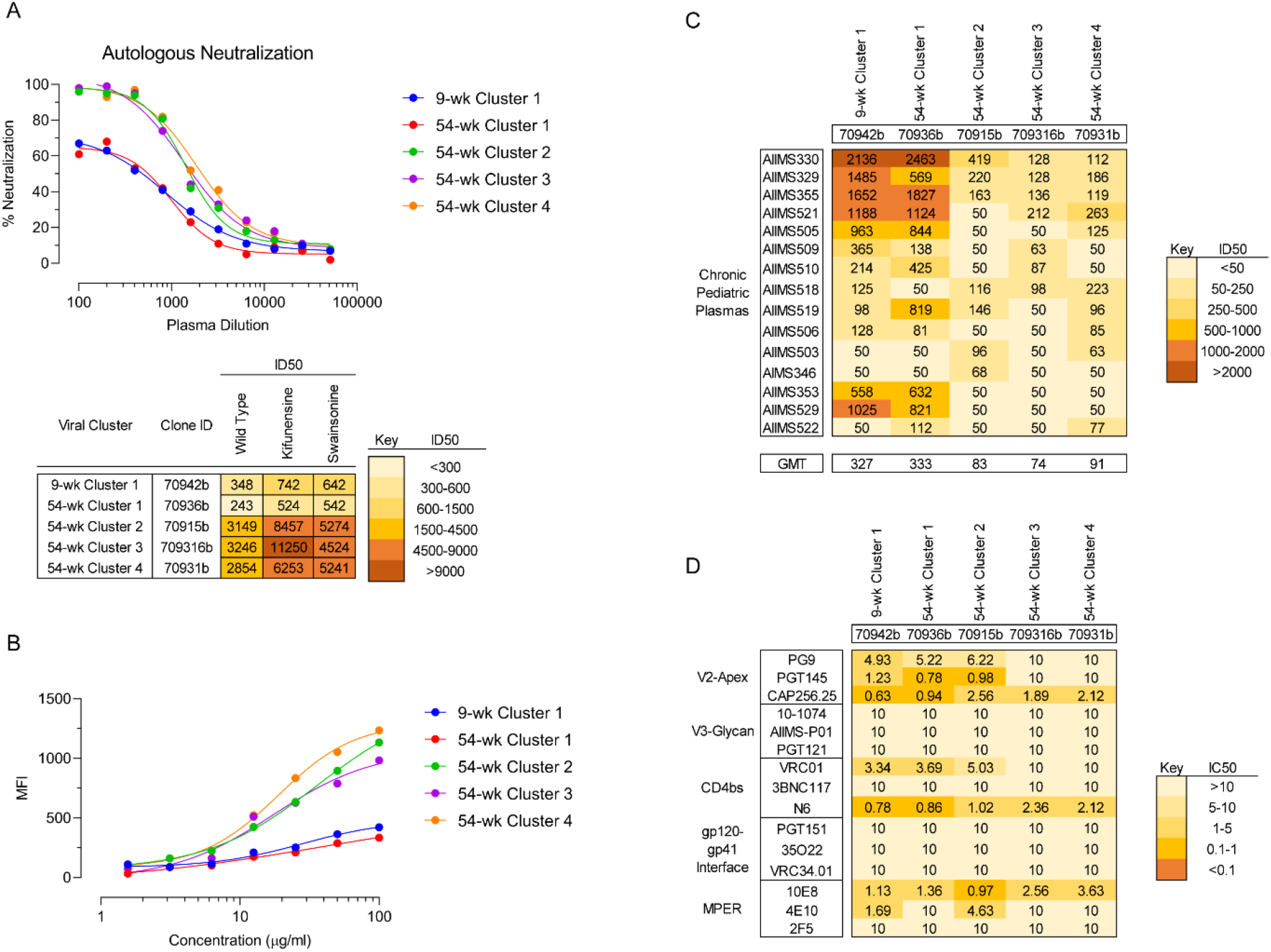
AIIMS709 plasma nAbs preferentially neutralized superinfecting viruses. (a) Neutralization curves of AIIMS709 viruses from 9-wk and 54-wk to autologous 54-wk plasma nAbs. ID_50_ values are shown for wildtype as well as hyperglycosylated variants (kifunensine and swainsonine treated) per the key given. (b) Surface binding assay of AIIMS709 9-wk and 54-wk viruses against 54-wk autologous IgG. (c) Neutralization titres of AIIMS709 9-wk and 54-wk viruses against heterologous plasma nAbs from chronic HIV-1 infected pediatric individuals with potent plasma nAbs. (d) Neutralization susceptibility of AIIMS709 9-wk and 54-wk viruses to known bnAbs targeting the V2-apex, V3-glycan, CD4 binding site (CD4bs), gp120-gp41 interface and/or fusion peptide, and membrane proximal external region (MPER). Shown are IC_50_ values. Neutralization assay were repeated thrice in triplicates and average values were used to draw curves and calculate ID_50_ and IC_50_ titres. Surface binding assay was repeated thrice and average MFI values were used for drawing curves. MFI – median fluorescence intensity.

The 54-wk cluster 2, 3 and 4 viral strains were the dominant circulating strains, while the 9-wk and 54-wk cluster 1 strains were primary infecting variants and present as minor variants in the 54-wk plasma (**figure 1**). The limited neutralization of the primary infecting variants by the 54-wk plasma nAbs as observed herein were contrary to our earlier observations of susceptibility of viruses from pediatric elite neutralizers to neutralization by the autologous plasma nAbs (5, 19). A plausible reason for the inability of the 54-wk plasma nAbs to effectively neutralize the 9-wk clade C strain, as compared to the contemporaneous A1C viruses, could be that the neutralization breadth induced by the superinfecting viruses is not caused by stimulation of the memory B cells developed in response to the primary infection but instead arose as de novo responses against the superinfecting strain and evolved with time, as was recently observed in superinfected women (20). A detailed analysis of the evolution of HIV-1 specific B cell responses during superinfection will enable understanding the mechanisms involved in the development of breadth of neutralizing antibody response. As the antibody responses during early HIV-1 infection are often non-neutralizing, it can be postulated that the circulating superinfecting A1C variants in this infant presented as immune complexes with the previously existing non-nAbs that were generated against the primary clade C variants. One of the factors contributing to the development of a relatively more potent and broader nAb response against superinfecting viruses herein is plausibly the formation of immune complexes, that are known to be more immunogenic than the antigen alone (21–23). Overall, the information acquired by evaluating the evolution of neutralizing antibody responses in a superinfected infant attribute the development of breadth of the AIIMS709 plasma nAb responses to the superinfecting A1C variants, targeted to the epitopes significantly exposed on A1C variants than primary infecting clade C variants. The findings provide an insight towards understanding the plausible mechanisms underlying the evolution of bnAbs during HIV-1 superinfection, in the setting of perinatal HIV-1 infection and remains to be further addressed in detail. Studies have reported that while the HIV-1 variants in early infection are resistant to neutralization by contemporaneous plasma antibodies, they become sensitive to plasma from later time points, indicating that de novo responses are subsequently generated against escape mutants, highlighting the dynamic nature of the neutralizing antibody response to HIV-1 (24, 25). In the case of this infant, plasma nAb response failed to show significant neutralization of early (primary) infecting variants, suggesting the distinctness of infant immune response to HIV-1as compared to adults.

To identify the exposed epitopes and neutralization phenotype of the four AIIMS709 viral variants (primary and superinfecting), we assessed their susceptibility to an exhaustive panel of bnAbs and non-nAbs targeting defined epitopes and plasma nAbs from chronically infected pediatric donors. In contrast to the neutralization susceptibility observed with autologous plasma nAbs, 54-wk cluster 2 and 3 variants were resistant, while 9-wk and 54-wk cluster 1 variants were sensitive to several plasma samples of chronically HIV-1 infected children (**figure 6C**), thereby highlighting the enhanced susceptibility of A1C superinfecting viruses to autologous plasma nAbs. Furthermore, the 54-wk cluster 2 and 3 viral strains (dominant circulating variants) were resistant to majority of known bnAbs except N6 (CD4bs targeting bnAb) and 10E8 (MPER targeting bnAb), whereas representative viral variants of cluster 1 and 3 were sensitive to most of the V2-glycan targeting bnAbs and CD4bs targeting bnAbs. All the viral variants in AIIMS709 were resistant to V3-glycan and interface targeting bnAbs (**figure 6D**). Sequence analysis indicated the absence of bnAb epitopes in A1C variants, corroborating with the observed resistance of these viruses to known bnAbs as well as pediatric plasmas with potent plasma bnAbs and further validating that the plasma nAbs in AIIMS709 targeted a distinct epitope, considering that the A1C variants were highly susceptible to 54-wk plasma nAbs.

We next generated several chimeras by swapping the 54-wk cluster 1 (clade C) gp120 regions into the 54-week cluster 3 viral strain (A1C), as it was the dominant circulating strain, to fine map the region targeted by autologous plasma nAbs on A1C viruses. The V1V2 and V5C5 swaps from the 54-wk cluster 1 into 54-wk cluster 3 rendered the viruses non-functional, while the C2 and C4 swaps did not show any variation in susceptibility profile. The C3V4 swaps showed a 10.7-fold reduction in ID_50_ titres (ID_50_ of 3246 vs 303), indicating that the plasma bnAbs targeted an epitope within the C3V4 region in cluster 3 viruses. Reverse swaps from the 54-wk cluster 3 into 54-wk cluster 1 variants increased susceptibility (4.2-fold increase, ID_50_ of 1020 vs 243) to plasma nAbs, further validating our results (**figure 7A – B**). Compared to the AIIMS709 clade C variants that had several glycan holes, the A1C variants had additional glycans in the C3V4 region (**figure 7C**). Though we were unable to decipher a dominant specificity by conventional epitope mapping, based on our observation of increased susceptibility of hyperglycosylated viral variants and computational prediction of genetic signatures within the C3V4 region responsible for neutralization by AIIMS709 plasma nAbs, we hypothesize these increased glycans in the autologous A1C variants were responsible for their unique neutralization phenotype. Previously, a longitudinal analysis of envelope sequences from the CAP-256 donor had revealed an overall increase in total PNGSs with oligomannose-type glycans during early infection, later followed by a decline in PNGSs due to loss of glycans at the V3 base and a decrease in oligomannose-type glycans, that was associated with the development of C3V4 directed nAbs. Based on the persistence of the intrinsic mannose patch over the course of HIV infection, it was proposed that this mannose rich region can serve as a stable HIV vaccine target (26). Further, it has been observed that the antibodies that target the high mannose patch are among the most potent and broadly HIV-1 nAbs known and are potential reagents for passive immunotherapy (4, 27).

**Figure 7.**
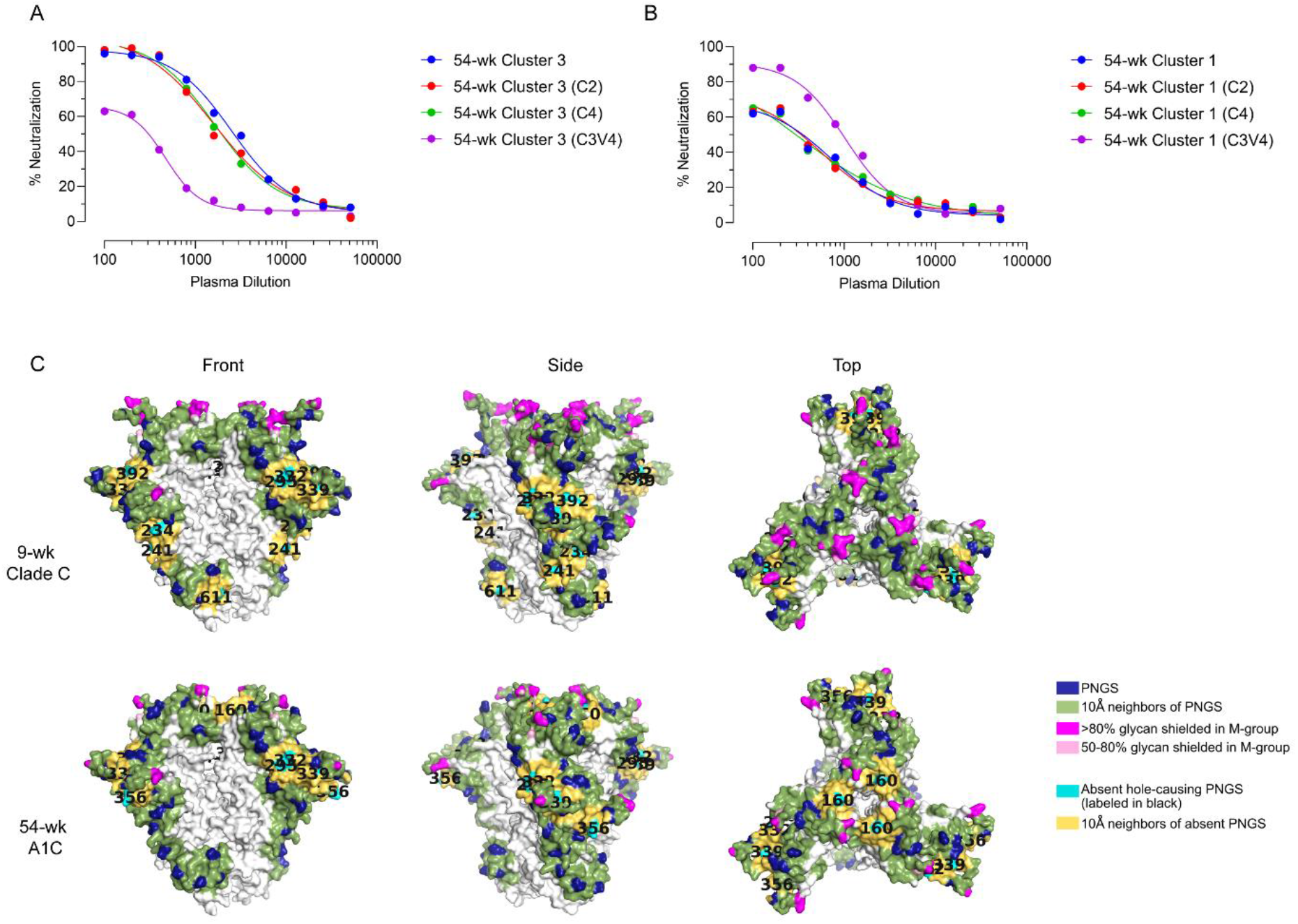
AIIMS709 plasma nAbs targeted glycan rich C3V4 region. (a – b) Neutralization curves of chimeric viruses generated by swapping C2, C3V4 and C4 region of 54-wk cluster 1 (clade C) and 54-wk cluster 3 (A1C) against autologous AIIMS709 54-wk plasma nAbs. Neutralization assay were repeated thrice in triplicates and average values were used to draw curves and calculate ID_50_ titres. (c) Predicted glycan shield for 9-wk cluster 1 (9-wk clade C) and 54-wk cluster 3 (54-wk A1C) showed prominent glycan holes in 9-wk clade C virus.

In summary, we report an HIV-1 superinfected infant in whom the nAb responses of unknown glycan specificities against the C3V4 region of the viral Env developed rapidly following superinfection and preferentially targeted superinfecting viral variants than primary infecting viral variants. Induction of nAbs effective against difficult-to-neutralize tier 2 HIV-1 variants will be a key aspect of a potential multistage HIV-1 vaccine. Herein we documented the impact of HIV-1 superinfection on evolution of the nAb responses in a perinatally infected infant. Our observation of rapid development of plasma nAbs that had a unique neutralization phenotype and preferentially neutralized superinfecting viruses than primary infecting viruses provides evidence that sequential immunization regimes can steer the development of potent bnAb responses. The observations herein are from a single case of superinfected infant. As HIV-1 superinfection in infants is a rare event, understanding the nAb responses in historical cases of HIV-1 superinfected infants can further provide mechanistic insight into how an immature immune system responds to the sequential exposure to distinct viral envelope glycoprotein (Env) towards the development of sequential immunization strategies to focus immune responses to conserved epitopes.

## Methods

### Study Subject

The current study was designed to address how HIV-1 superinfection in an infant would drive bnAb responses, in the setting of an immature immune system. After written informed consent from the guardians of AIIMS709, blood was drawn from two time-points (corresponding to 9-week and 54-week post-infection) in 3-ml EDTA vials, plasma was aliquoted for plasma neutralization assays, viral RNA isolation, and viral loads. The study was approved by institute ethics committee of All India Institute of Medical Sciences (IECPG-307/07.09.2017).

### Plasmids, viruses, monoclonal antibodies, and cells

Plasmids encoding HIV-1 env genes representing different clades, monoclonal antibodies and TZM-bl cells were procured from NIH AIDS Reagent Program. 10-1074 and BG18 expression plasmids were kindly provided by Dr. Michel Nussenzweig, Rockefeller University, USA. CAP256.09, CAP256.25 and b6 were procured from IAVI Neutralizing Antibody Centre, USA. HEK293T cells were purchased from the American Type Culture Collection (ATCC).

### HIV-1 envelope sequences and phylogenetic analysis

HIV-1 envelope genes were PCR amplified from AIIMS709 9-wk and 54-wk plasma viral RNA by single genome amplification and directly sequenced commercially. Individual sequence fragments of SGA amplified amplicons were assembled using Sequencher 5.4 (Gene Code Corporation). Subtyping for SGA sequences was performed with REGA HIV subtyping tool (400bp sliding window with 200 bp steps size). Inter-clade recombination was examined with RIP 3.0 (Recombinant Identification Program). Nucleotide sequences were aligned with MUSCLE in MEGA X 10.1. Maximum-likelihood trees were computed with MEGA X 10.1 using a general-time reversal substitution model incorporating a discrete gamma distribution with 5 invariant sites. Evolutionary divergence within each infant’s SGA sequence was conducted in MEGA X 10.1 and was calculated as number of base substitutions per site from averaging over all sequence pairs. Analyses were conducted using the Maximum Composite Likelihood model. The rate variation among sites was modelled with a gamma distribution (shape parameter = 5).

### Generation of replication incompetent wildtype and chimeric pseudoviruses

Autologous replication incompetent envelope pseudoviruses were generated from AIIMS709’s 9-wk and 54-wk plasma sample. Viral RNA was isolated from 140 μl of plasma using QIAamp Viral RNA Mini Kit, reverse transcribed, using gene specific primer OFM19 (5’ - GCACTCAAGGCAAGCTTTATTGAGGCTTA – 3’) and Superscript III reverse transcriptase, into cDNA which was used in two-round nested PCR for amplification of envelope gene using High Fidelity Phusion DNA Polymerase (New England Biolabs). First round primers consisted of forward primer VIF2 (5’ – GGGTTTATTACAGAGACAGCAGAG – 3’) and reverse primer OFM19 (5’ – GCACTCAAGGCAAGCTTTATTGAGGCTTA – 3’). Second round primers consisted of forward primer ENVA (5’ - CACCGGCTTAGGAATTTACTATGGCAGGAAG - 3’) and reverse primer ENVN (5’ - TGCCAATCAGGGAAAAAGCCTTGTGTG - 3’. The envelope amplicons were purified, and ligated into pcDNA3.1D/V5-His-TOPO vector (Invitrogen). Chimeric envelope pseudoviruses were generated by swapping the V1V2, C2, C3V4, C4, V5C5 C5 regions between 9-wk and 54-wk viruses by individually amplifying the respective regions via conventional PCR and the remaining backbone by inverse PCR. The amplified regions (V1V2, C2, C3V4, C4, V5C5 C5) were then ligated into the backbone with infusion HD cloning plus kit as per manufacturer’s instructions.

Pseudoviruses were prepared by co-transfecting 1.25 μg of HIV-1 envelope containing plasmid with 2.5 μg of an envelope deficient HIV-1 backbone (PSG3Δenv) vector at a molar ratio of 1:2 using PEI-MAX as transfection reagent in HEK293T cells seeded in a 6-well culture plates. For generation of hyperglycosylated pseudoviruses, one hour prior to transfection, HEK293T cells were treated with either Kifunensine (25 μM) or Swainsonine (20 μM). Culture supernatants containing pseudoviruses were harvested 48 hours post-transfection, filtered through 0.4μ filter, aliquoted and stored at −80°C until further use. TCID_50_ was determined by infecting TZM-bl cells with serially diluted pseudoviruses in presence of DEAE-Dextran, and lysing the cells 48 hours post-infection. Infectivity titres were determined by measuring luminescence activity in presence of Bright Glow reagent (Promega).

### Neutralization assay

Neutralization assays were carried out using TZM-bl cells, a genetically engineered HeLa cell line that constitutively expresses CD4, CCR5 and CXCR4, and contains luciferase and β-galactosidase gene under HIV-1 tat promoter. Envelope pseudoviruses were incubated in presence of serially diluted heat inactivated plasmas, bnAbs or non-nAbs for one hour. After incubation, freshly Trypsinized TZM-bl cells were added, with 25 μg/ml DEAE-Dextran. The plates were incubated for 48h at 37°C, cells were lysed in presence of Bright Glow reagent, and luminescence was measured. Using the luminescence of serially diluted bnAbs or plasma, a non-linear regression curve was generated and titres were calculated as the bnAb concentration, or reciprocal dilution of serum that showed 50% reduction in luminescence compared to untreated virus control.

### Cell surface binding assay

1.25 x 10^5^ HEK293T cells seeded in a 12-well plate were transiently transfected with 1.25 μg of env-coding plasmids (pcDNA3.1 with cloned env/rev cassettes) using PEI-MAX. 48 hours post-transfection, cells were harvested and per experimental requirement, distributed in 1.5 ml microcentrifuge tubes. For monoclonal antibody staining, 10 μg/ml of antibody was used and titrated 2-fold in staining buffer. 100 μl of primary antibody (HIV-1 specific monoclonals) were added to HEK293T cells expressing envs, and incubated for 30 minutes at room temperature. After washing, 100 μl of 1:500 diluted PE conjugated mouse anti-human IgG Fc was added, and after 30-minute incubation, a total of 50,000 cells were acquired on BD LSRFortessa X20. Data was analyzed using FlowJo software (version v10.6.1).

### Statistical analysis

2-tailed student’s t test for paired analysis and Mann-Whitney U test for unpaired analysis were used. For assessing the relative infectivity, area under curves were calculated. All statistical analyses were performed on GraphPad Prism 8. A p-value of <0.05 was considered significant. Neutralization assays were performed in triplicates and repeated thrice. Average ID50 values are shown and used for statistical comparisons. Binding ELISAs were performed in duplicates and repeated thrice. Average OD450 values were used for plotting curves. Surface binding assay was performed thrice and average PE-MFI (phycoerythrin-median fluorescence intensity) values were used for plotting curves.

## Acknowledgments

We thank all the study subjects for participating in this study. We are thankful to NIH AIDS Reagent program for providing HIV-1 envelope pseudovirus plasmids, bnAbs, non-nAbs and their expression plasmids, and TZM-bl cells, and Neutralizing Antibody Consortium (NAC), IAVI, USA for providing bnAbs. We are thankful to Dr. Michel Nussenzweig for providing 10-1074 and BG18 bnAb expression plasmids.

## Author contributions

N.M designed the study, performed SGA amplification, pseudovirus cloning, and neutralization assays, analyzed data, wrote the initial manuscript, revised and finalized the manuscript. S.S, and A. D contributed to SGA amplification, pseudovirus cloning, and neutralization assays. S.K, H.C and Sw.S expressed PGT145, PGDM1400, CAP256.25, BG18, 10-1074 and AIIMS-P01 bnAb. R.S, and B. K.D provided the immunological data of the HIV-1 infected infants. R.L, and Su.K.K, provided the samples of HIV-1 infected infants and provided patient care and management. S.S, A.D, S.K, and H.C edited and revised the manuscript. K.L conceptualized and designed the study, edited, revised and finalized the manuscript.

## Data Availability

The SGA amplified HIV-1 envelope sequences used for inference of phylogeny and highlighter plots are available at GenBank with accession numbers MN703360 – MN703366. All data required to state the conclusions in the paper are present in the paper and/or the supplementary data. Source data are provided with this paper. Additional information related to the paper, if required, can be requested from the authors.

## References

1. Bonsignori M, Liao H-X, Gao F, Williams WB, Alam SM, Montefiori DC, Haynes BF. 2017. Antibody-virus co-evolution in HIV infection: paths for HIV vaccine development. Immunol Rev 275:145–160.

2. Fouda GG, De Paris K, Levy O, Marchant A, Gray G, Permar S, Marovich M, Singh A. 2020. Immunological mechanisms of inducing HIV immunity in infants. Vaccine 38:411–415.

3. Goo L, Chohan V, Nduati R, Overbaugh J. 2014. Early development of broadly neutralizing antibodies in HIV-1-infected infants. Nat Med 20:655–658.

4. Kwong PD, Mascola JR. 2018. HIV-1 Vaccines Based on Antibody Identification, B Cell Ontogeny, and Epitope Structure. Immunity 48:855–871.

5. Mishra N, Sharma S, Dobhal A, Kumar S, Chawla H, Singh R, Makhdoomi MA, Das BK, Lodha R, Kabra SK, Luthra K. 2020. Broadly neutralizing plasma antibodies effective against autologous circulating viruses in infants with multivariant HIV-1 infection. Nat Commun 11:4409.

6. Sok D, Burton DR. 2018. Recent progress in broadly neutralizing antibodies to HIV. Nat Immunol 19:1179–1188.

7. Kumar S, Panda H, Makhdoomi MA, Mishra N, Safdari HA, Chawla H, Aggarwal H, Reddy ES, Lodha R, Kumar Kabra S, Chandele A, Dutta S, Luthra K. 2018. An HIV-1 broadly neutralizing antibody from a clade C infected pediatric elite neutralizer potently neutralizes the contemporaneous and autologous evolving viruses. J Virol https://doi.org/10.1128/JVI.01495-18.

8. Simonich CA, Williams KL, Verkerke HP, Williams JA, Nduati R, Lee KK, Overbaugh J. 2016. HIV-1 Neutralizing Antibodies with Limited Hypermutation from an Infant. Cell 166:77–87.

9. Muenchhoff M, Adland E, Karimanzira O, Crowther C, Pace M, Csala A, Leitman E, Moonsamy A, McGregor C, Hurst J, Groll A, Mori M, Sinmyee S, Thobakgale C, Tudor-Williams G, Prendergast AJ, Kloverpris H, Roider J, Leslie A, Shingadia D, Brits T, Daniels S, Frater J, Willberg CB, Walker BD, Ndung’u T, Jooste P, Moore PL, Morris L, Goulder P. 2016. Nonprogressing HIV-infected children share fundamental immunological features of nonpathogenic SIV infection. Sci Transl Med 8:358ra125.

10. Hraber P, Korber B, Wagh K, Montefiori D, Roederer M. 2018. A single, continuous metric to define tiered serum neutralization potency against HIV. Elife 7.

11. Scanlan CN, Offer J, Zitzmann N, Dwek RA. 2007. Exploiting the defensive sugars of HIV-1 for drug and vaccine design. Nature 446:1038–1045.

12. Andrabi R, Voss JE, Liang C-H, Briney B, McCoy LE, Wu C-Y, Wong C-H, Poignard P, Burton DR. 2015. Identification of Common Features in Prototype Broadly Neutralizing Antibodies to HIV Envelope V2 Apex to Facilitate Vaccine Design. Immunity 43:959–973.

13. Andrabi R, Su C-Y, Liang C-H, Shivatare SS, Briney B, Voss JE, Nawazi SK, Wu C-Y, Wong C-H, Burton DR. 2017. Glycans Function as Anchors for Antibodies and Help Drive HIV Broadly Neutralizing Antibody Development. Immunity 47:524–537.e3.

14. Crooks ET, Grimley SL, Cully M, Osawa K, Dekkers G, Saunders K, Rämisch S, Menis S, Schief WR, Doria-Rose N, Haynes B, Murrell B, Cale EM, Pegu A, Mascola JR, Vidarsson G, Binley JM. 2018. Glycoengineering HIV-1 Env creates “supercharged” and “hybrid” glycans to increase neutralizing antibody potency, breadth and saturation. PLoS Pathog 14:e1007024.

15. Landais E, Huang X, Havenar-Daughton C, Murrell B, Price MA, Wickramasinghe L, Ramos A, Bian CB, Simek M, Allen S, Karita E, Kilembe W, Lakhi S, Inambao M, Kamali A, Sanders EJ, Anzala O, Edward V, Bekker L-G, Tang J, Gilmour J, Kosakovsky-Pond SL, Phung P, Wrin T, Crotty S, Godzik A, Poignard P. 2016. Broadly Neutralizing Antibody Responses in a Large Longitudinal Sub-Saharan HIV Primary Infection Cohort. PLoS Pathog 12:e1005369.

16. Lei L, Yang YR, Tran K, Wang Y, Chiang C-I, Ozorowski G, Xiao Y, Ward AB, Wyatt RT, Li Y. 2019. The HIV-1 Envelope Glycoprotein C3/V4 Region Defines a Prevalent Neutralization Epitope following Immunization. Cell Rep 27:586–598.e6.

17. Moore PL, Gray ES, Sheward D, Madiga M, Ranchobe N, Lai Z, Honnen WJ, Nonyane M, Tumba N, Hermanus T, Sibeko S, Mlisana K, Abdool Karim SS, Williamson C, Pinter A, Morris L, CAPRISA 002 Study. 2011. Potent and broad neutralization of HIV-1 subtype C by plasma antibodies targeting a quaternary epitope including residues in the V2 loop. J Virol 85:3128–3141.

18. Zhao F, Joyce C, Burns A, Nogal B, Cottrell CA, Ramos A, Biddle T, Pauthner M, Nedellec R, Qureshi H, Mason R, Landais E, Briney B, Ward AB, Burton DR, Sok D. 2020. Mapping Neutralizing Antibody Epitope Specificities to an HIV Env Trimer in Immunized and in Infected Rhesus Macaques. Cell Rep 32:108122.

19. Mishra N, Makhdoomi MA, Sharma S, Kumar S, Kumar D, Chawla H, Singh R, Kanga U, Das BK, Lodha R, Kabra S, Luthra K. 2018. Viral characteristics associated with sustenance of elite neutralizing activity in chronically HIV-1C infected monozygotic pediatric twins. bioRxiv 475822.

20. Sheward DJ, Marais J, Bekker V, Murrell B, Eren K, Bhiman JN, Nonyane M, Garrett N, Woodman ZL, Abdool Karim Q, Abdool Karim SS, Morris L, Moore PL, Williamson C. 2018. HIV Superinfection Drives De Novo Antibody Responses and Not Neutralization Breadth. Cell Host Microbe 24:593–599.e3.

21. Brady LJ. 2005. Antibody-mediated immunomodulation: a strategy to improve host responses against microbial antigens. Infect Immun 73:671–678.

22. Desikan R, Raja R, Dixit NM. 2020. Early exposure to broadly neutralizing antibodies may trigger a dynamical switch from progressive disease to lasting control of SHIV infection. PLoS Comput Biol 16:e1008064.

23. Shubin Z, Li W, Poonia B, Ferrari G, LaBranche C, Montefiori D, Zhu X, Pauza CD. 2017. An HIV Envelope gp120-Fc Fusion Protein Elicits Effector Antibody Responses in Rhesus Macaques. Clin Vaccine Immunol 24.

24. Anthony C, York T, Bekker V, Matten D, Selhorst P, Ferreria R-C, Garrett NJ, Karim SSA, Morris L, Wood NT, Moore PL, Williamson C. 2017. Cooperation between Strain-Specific and Broadly Neutralizing Responses Limited Viral Escape and Prolonged the Exposure of the Broadly Neutralizing Epitope. J Virol 91.

25. Sather DN, Carbonetti S, Malherbe DC, Pissani F, Stuart AB, Hessell AJ, Gray MD, Mikell I, Kalams SA, Haigwood NL, Stamatatos L. 2014. Emergence of broadly neutralizing antibodies and viral coevolution in two subjects during the early stages of infection with human immunodeficiency virus type 1. J Virol 88:12968–12981.

26. Coss KP, Vasiljevic S, Pritchard LK, Krumm SA, Glaze M, Madzorera S, Moore PL, Crispin M, Doores KJ. 2016. HIV-1 Glycan Density Drives the Persistence of the Mannose Patch within an Infected Individual. J Virol 90:11132–11144.

27. Daniels CN, Saunders KO. 2019. Antibody responses to the HIV-1 envelope high mannose patch. Adv Immunol 143:11–73.

